# The genome of the Balearic shearwater (*Puffinus mauretanicus*), a Critically Endangered seabird: a valuable resource for evolutionary and conservation genomics

**DOI:** 10.1101/2021.12.17.473171

**Authors:** Cristian Cuevas-Caballé, Joan Ferrer Obiol, Joel Vizueta, Meritxell Genovart, Jacob Gonzalez-Solís, Marta Riutort, Julio Rozas

## Abstract

The Balearic shearwater (*Puffinus mauretanicus*) is the most threatened seabird in Europe. The fossil record suggests that human colonisation of the Balearic Islands resulted in a sharp decrease of the population size. Currently, populations continue to be decimated mainly due to predation by introduced mammals and bycatch in longline fisheries, and some studies predict their extinction by 2070. We present the first high-quality reference genome for the species which was obtained by a combination of short and long-read sequencing. Our hybrid assembly includes 4,169 scaffolds, with a scaffold N50 of 2.1 Mbp, a genome length of 1.2 Gbp, and BUSCO completeness of 96%, which is amongst the highest across sequenced avian species. This reference genome allowed us to study critical aspects relevant to the conservation status of the species, such as an evaluation of overall heterozygosity levels and the reconstruction of its historical demography. Our phylogenetic analysis using whole-genome information resolves current uncertainties in the order Procellariiformes systematics. Comparative genomics analyses uncover a set of candidate genes that may have played an important role into the adaptation to a pelagic lifestyle of Procellariiformes, including those for the enhancement of fishing capabilities, night vision and the development of natriuresis. This reference genome will be the keystone for future developments of genetic tools in conservation efforts for this Critically Endangered species.

## Background

The genomic sequence of a species accumulates valuable information on the evolutionary history, including demographic and selective events, and on the evolution of genes and traits [1–4], information that it is also crucial for the emerging field of conservation genomics [5]. The genetic diversity within a species represents a reservoir of adaptive variation that can help populations to cope with environmental variability [6]. Understanding the processes that shape genetic diversity and its distribution pattern within species is paramount to assess the conservation status or the factors responsible for a species decline [7] [8]. This knowledge can inform the proposal of effective conservation and management plans, as for instance the definition of management units [9]. In this context, next-generation sequencing (NGS) techniques allow the analysis of an increased density of markers across the genome, providing unprecedented accuracy in the estimations of population genetic parameters relevant for scientific-based conservation recommendations [10].

Among the Critically Endangered species listed by the IUCN Red List (IUCN 2021), we find the Balearic shearwater (*Puffinus mauretanicus* Lowe, 1921) (Figure 1a) belonging to the most diverse order of seabirds, the Procellariiformes. This order has a worldwide distribution and comprises more than 140 species (IUCN 2021) in four families: petrels and shearwaters (Procellariidae); northern storm petrels (Hydrobatidae); southern storm petrels (Oceanitidae) and albatrosses (Diomedeidae). All species show many morphological, physiological and life history traits associated with their adaptation to a pelagic lifestyle. They are long-lived with deferred sexual maturity, low fecundity (all lay a single egg), colonial breeders, socially (and mostly sexually) monogamous, highly phylopatric and with prominent salt gland at the base of the bill, adaptations for underwater vision to fish as well as a particularly acute sense of smell, among other traits [12]. Within this apparent homogeneity, the group shows a large variation in body mass and lifestyles, ranging from 20g to 15kg, from bodies shaped for diving (e.g. short strong wings used for wing-propelled diving) to those prepared for an extremely vagile lifestyle (thin elongated wings) and from a continuous flapping to dynamic soaring flight modes. Currently, their phylogenetic relationships present some conflicting issues [13,14], such as the position of albatrosses, whether storm petrels (Families Hydrobatidae and Oceanitidae) constitute a monophyletic group, and whether diving-petrels (genus *Pelecanoides*) should be considered an independent family from Procellariidae.

**Figure 1.**
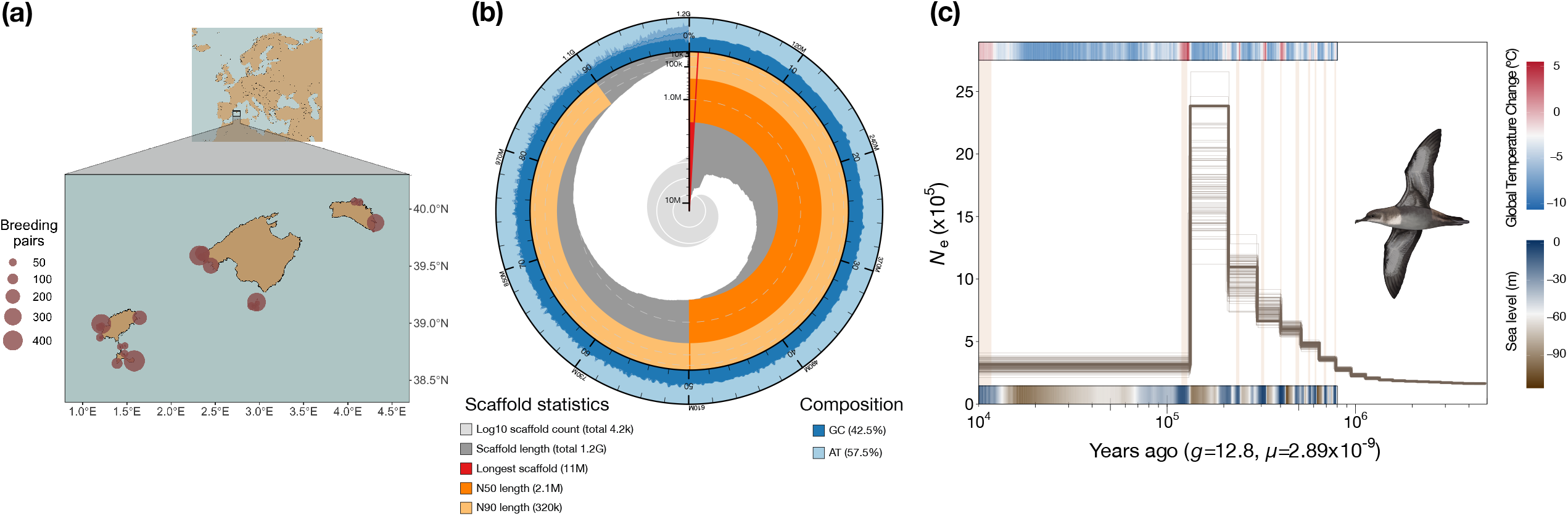
(a) Map depicting the known Balearic shearwater breeding colonies in the Balearic Islands. Circle size is proportional to population size as shown in the legend. Modified from [20] (b) Snail plot summarizing genome assembly statistics [120]. From inside to outside, the light-grey spiral shows the cumulative scaffold count on a log scale with white scale lines depicting changes of order of magnitude. Dark-grey segments show the distribution of scaffold lengths, and the plot radius is scaled to the longest scaffold (shown in red). Orange and light-orange rings represent the N50 and N90 scaffold lengths, respectively. Blue and light-blue rings show GC, AT and N percentages along the genome assembly. (c) MSMC2 reconstruction of effective population size estimates (*N*_e_) over time, estimated using generation time of 12.8 years and mutation rate of 2.89 × 10^−9^ substitutions per nucleotide per generation. Light-brown vertical bars represent interglacial periods. Upper panel represents global temperature changes as inferred from the EPICA (European Project for Ice Coring in Antarctica) Dome C ice core [121]. Lower panel represents sea level changes inferred from a stack of 57 globally distributed benthic δ 18O records [122].

The Balearic shearwater is a medium-sized pelagic seabird endemic to the Balearic Islands. Its population size is undergoing a fast annual decline of 7.4-14% [15,16] mostly due to bycatch in longline fisheries and predation by invasive mammals in the colonies [17–19]. Currently, it has a reduced number of breeding pairs (estimated as *ca*. 3,200, Arcos, 2011, with a total population size up to 30,000 individuals due to the vast contingent of floaters [21,22]. Genetic studies based on mtDNA and microsatellites found that this species has low levels of genetic diversity and high inbreeding coefficients [23]. Although local inbreeding and natal philopatry can contribute to a reduction in population size, the actual worst menace for the species comes from human activities, and a population viability study based on demographic modeling predicted that the species would become extinct by 2070 [15].

Indeed, a population viability study based on demographic modeling predicted that the species would become extinct by 2070 [15]. Moreover, studies based on mitochondrial markers [24] and also on morphology and migratory behaviour [25], suggested a possible ongoing hybridization and introgression process between Balearic and Mediterranean (*P. yelkouan*) shearwaters, which may represent an additional threat for the species.

Here, we (1) provide a high-quality reference genome for the Balearic shearwater along its structural and functional annotations; (2) estimate genome-wide heterozygosity and the historical demography of the species by performing Multiple Sequentially Markovian Coalescent (MSMC) analyses; (3) revisit the phylogeny of the order by using this genome together with seven additional Procellariiformes genomes released by the B10K [3], and (4) uncover genes putatively involved in Procellariiformes adaptation to pelagic life. The high-quality genome of the most endangered seabird in Europe presented here will be the base for further population-based conservation genomics studies.

### Data description

#### Sampling, DNA and RNA extraction and sequencing

We sampled two Balearic shearwater adults and one chick. Adults were sampled on Sa Cella colony, Mallorca (male) and on Sa Conillera, Eivissa (unsexed) in 2004, while the chick was sampled on Conills islet (Mallorca) in July 2019. From here on the animals will be referred to as male-Mll, unsexed-Ei and chick-Mll, respectively. Special permits to obtain the samples were issued by Conselleria de Medi Ambient, Agricultura i Pesca (Govern de les Illes Balears, Spain).

We extracted DNA from blood samples preserved in absolute ethanol for both adults. The DNA extraction for the male-Mll was performed with DNeasy Blood & Tissue Kit (Qiagen) following the manufacturer’s instructions, and with Blood & Cell Culture DNA Mini Kit (Qiagen) for the unsexed-Ei. RNA was extracted from the chick-Mll’s blood cells preserved in RNAlater 1:5 using the RNeasy Mini Kit (Qiagen) according to the manufacturer’s protocols. We performed the quality control with gel electrophoresis and NanoDrop One (Thermo Fisher Scientific, Waltham, MA, USA), and the quantification with an Invitrogen Qubit Fluorometer 2.0 (Broad Range kit).

We obtained the reference genome combining short-read and long-read sequencing libraries, and using RNA-seq data to assist with the annotation. First, an Illumina TruSeq DNA PCR Free library (insert size = 350 bp) was prepared by Macrogen (South Korea) using DNA from male-Mll, and sequenced using two HiSeq X Ten runs (2×150bp). Second, long-read libraries were prepared, from the DNA of unsexed-Ei, using the Ligation kit SQK-LSK109 1D from ONT (Oxford Nanopore Technologies) (N50 of 9431 bp) at CNAG (Centro Nacional de Análisis Genómico, Spain) and sequenced through five runs of MinION on FLO-MIN106 flow cells. Third, we prepared RNA sequencing (RNA-seq) libraries from the chick-Mll’s RNA using the TruSeq RNA Sample Prep Kit v2 with Ribo-Zero, and we sequenced the libraries on a NovaSeq 6000 (2×100bp) (Macrogen, South Korea).

### Analyses

#### Sequencing data and Genome assembly and annotation

Illumina paired-end (2×150 bp) sequencing of the male-Mll yielded a throughput of 147.7 Gbp (Table 1), representing a mean coverage of 118x. The five runs of ONT sequencing of the unsexed-Ei resulted in a 10x coverage with a read N50 of 9,431 bp. RNA sequencing of the chick-Mll (2×100 bp) yielded 15 Gbp of data.

**Table 1.** Sequencing data, library information and samples used in this study.

| Library | Total number of base pairs | Number of reads | Coverage | Individual code | Location |
| --- | --- | --- | --- | --- | --- |
| HiSeq X Ten - TruSeq DNA PCR Free. 2x150 bp | 143,765,593,200 | 958,437,288 | 118x | Male-MII | Sa Cella (Mallorca) |
| ONT - Ligation kit SQK-LSK109 1D | 12,142,789,693 | 2,576,486 | 10x | Unsexed-Ei | Sa Conillera (Eivissa) |
| NovaSeq 6000 - TruSeq RNA Sample Prep Kit v2. 2x100 bp | 14,997,592,000 | 149,975,920 | -- | Chick-MII | Conills islet (Mallorca) |

We obtained a hybrid assembly with MaSuRCA formed by 4,169 scaffolds, with an N50 of 2.1 Mbp, and an assembly length of 1.21 Gbp (Table 2, Figure 1b). The completeness analysis using BUSCO yields a value of 95.9%, and only 0.3% of the complete genes were duplicated and 1.1% were fragmented (Table 2). Our *de novo* repeat annotation analysis shows that 9.95% of the genome consists of repetitive regions (Table S1), which is within the range of previously sequenced avian genomes [26]. Among repeat elements, long interspersed nuclear elements (LINEs) were the most abundant (4.45% of the genome). The genome annotation process resulted in a total of 21,959 protein-coding genes, of which 18,769 (85.5%) have at least one GO associated term, and 19,218 (87.5%) have hits across the surveyed curated databases (Table S2).

**Table 2.** Balearic shearwater genome assembly metrics.

|  |  |
| --- | --- |
| Assembly length (bp) | 1,218,519,395 |
| Number of scaffolds | 4,169 |
| Longest scaffold (Mbp) | 11.06 |
| N50 (Mbp) | 2.13 |
| L50 | 164 |
| GC content (%) | 42.52 |
| Repetitive content (%) | 9.95 |
| Mitogenome (bp) | 19,855 |
| No. protein-coding genes | 21,959 |
| BUSCO % |  |
| Complete | 95.9 |
| Single copy | 95.6 |
| Duplicated | 0.3 |
| Fragmented | 1.1 |
| Missing | 3.0 |

Blood transcriptome assembly from the chick-Mll resulted in 224,904 transcripts (Table S3). However, BUSCO completeness was only 62.4%, which was far below genome completeness, probably due to the RNA coming from a single not very transcriptionally active tissue.

The assembly of the mitogenome of *P. mauretanicus* resulted in a single contig of 19,885 bp long, with a coverage (Illumina reads) of 371x, which is around three times higher than the coverage of the nuclear genome. This mitogenome has the same gene order as other published Procellariiformes’ mitogenomes (Figure S1). The mitogenome has two copies of the *nad6* gene, as predicted in *P. lherminieri* [27]; the later feature was also confirmed analysing the mean coverage (illumina reads) across genes (Table S4).

#### Historical demography of the Balearic shearwater

Balearich shearwater PSMC’ analysis showed support for a steady growth in population size from an originally low population size followed by a sudden increase ∼200 kya (Figure 1c). High population size did not last long and suffered a sudden decrease to nearly one tenth of the population coinciding with the end of the glacial period before the last interglacial period (119-128 kya) and a prolonged period of low sea level (Figure 1c). Hereafter, *N*_e_ remained stable until ∼10 kya ago, as more recent MSMC2 time segments are regarded as being unreliable [28].

Genome-wide heterozygosity in *P. mauretanicus* was 0.0024, which is within the range of genome-wide heterozygosities estimated for other Procellariiformes (ranging from 0.0014 in *T. chlororhynchos* to 0.0037 in *P. urinatrix)* (Figure 2a). Among Procellariiformes, small-bodied species tended to have higher mean heterozygosities but also higher variance than large-bodied species (Figure 2b).

**Figure 2.**
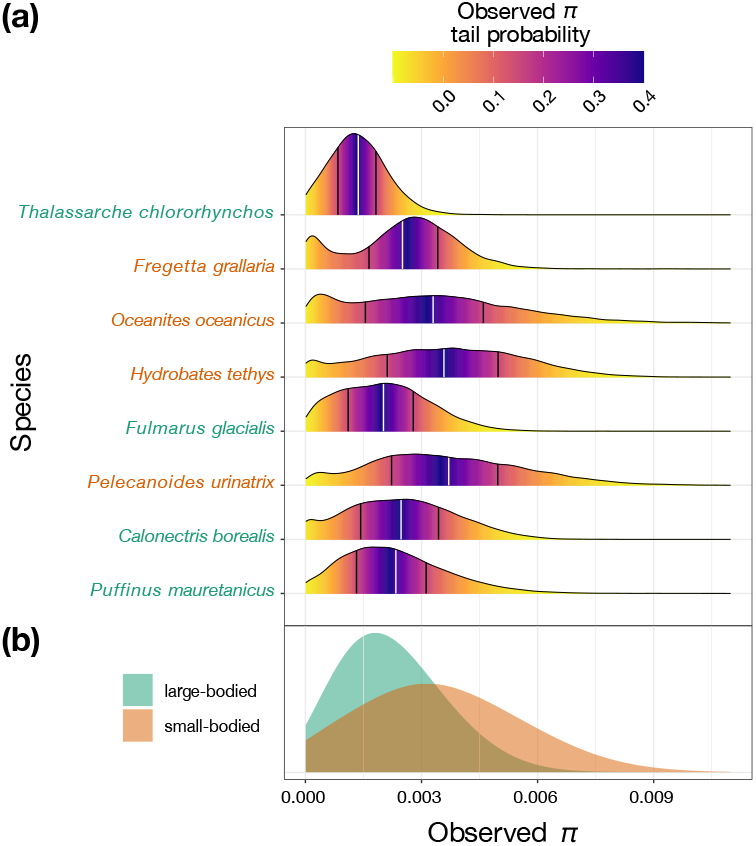
Comparison of genome-wide heterozygosity among Procellariiformes. (a) Density plots showing the distribution of individual nucleotide diversity (*π*) values in nonoverlapping 25Kb windows for each of the eight Procellariiformes species with an available reference genome. Scientific names of large-bodied and small-bodied species are shown in green and orange, respectively. Color-scale represents *π* values tail probabilities as shown in the legend. The white line depicts median values and black lines depict 25th and 75th percentiles. (b) Density plots showing the distribution of *π* values in large-bodied and small-bodied species groups.

#### Phylogenetic relationships

OrthoFinder analysis estimated 6,172 single copy (1:1) ortholog genes across the 12 genomes surveyed. With this data we generated three supermatrices: 1) CDS supermatrix of 10,534,506 bp long to extract the 4D sites, 2) 4D supermatrix with 1,512,677 4-fold degenerate sites, and 3) the amino acid supermatrix including 3,466,564 sites. Phylogenetic analyses using the 4D and the amino acid supermatrices recovered the same topology with full support at all nodes (ultrafast bootstrap = 100; Figures S2 and S3).

The analysis performed to explicitly account for incomplete lineage sorting (ILS) with ASTRAL using either the individual gene sequences (CDS gene trees) or the individual amino acid sequences (amino acid gene trees), produced species trees with the same topology as those obtained by ML using 4D or amino acid supermatrices (Figures S4 and Figure S5). The normalized quartet score (proportion of input gene tree quartet trees in agreement with the species tree) was 0.78 for CDS gene trees and 0.64 for amino acid gene trees.

The ultrametric tree (Figure 3) obtained using r8s from the 4D supermatrix ML tree summarizes the recovered topology. In this topology, the Atlantic yellow-nosed albatross (*T. chlororhynchos*, Diomedeidae) is the sister group to all the other Procellariiformes. We also find that storm petrels (Hydrobatidae and Oceanitidae) do not constitute a monophyletic group. In addition, diving petrels (*Pelecanoides*) are included within Procellariidae.

**Figure 3.**
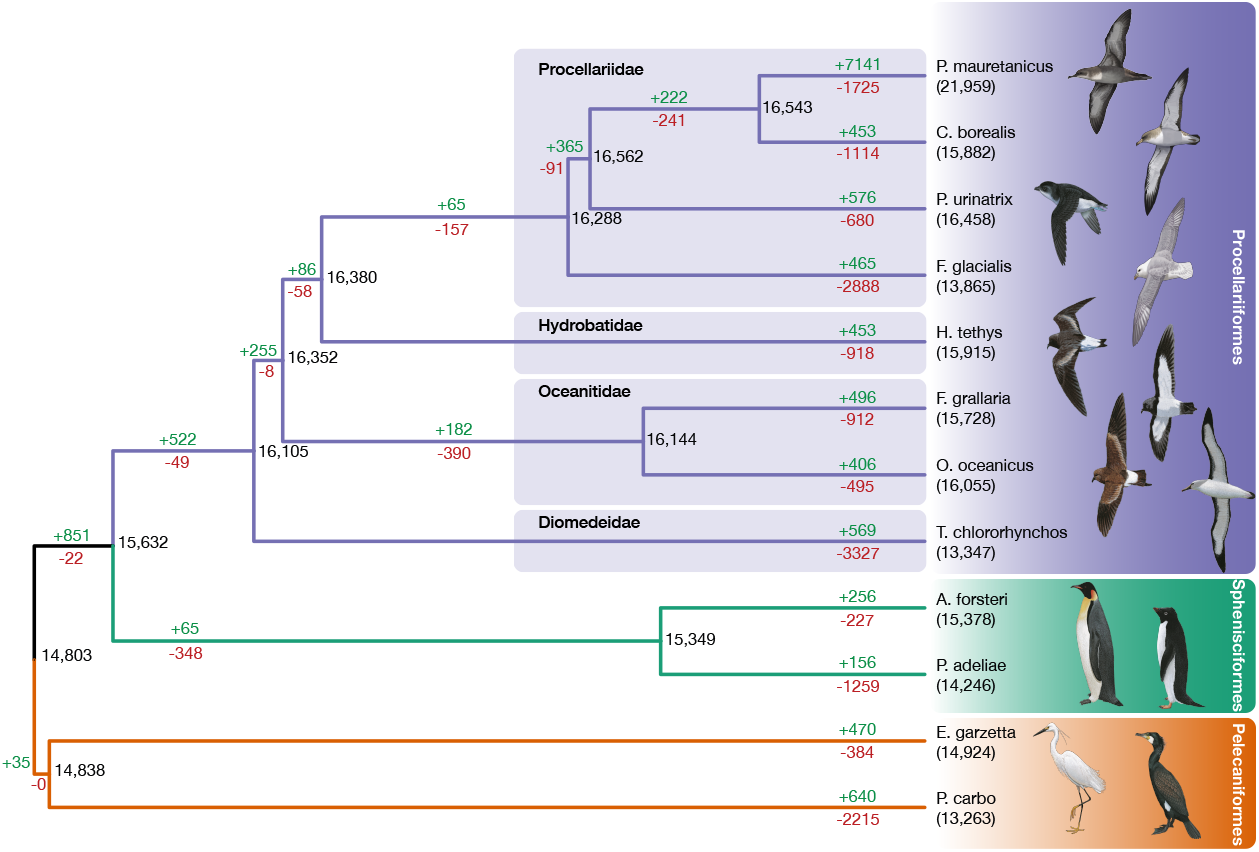
Ultrametric tree based on the 4D CDS ML tree calibrated with r8s. Minimum number of gains (green) and losses (red) per branch are represented according to BadiRate analysis. Numbers in ancestral nodes and in the tips (in parenthesis) indicate the inferred number of genes. Illustrations of seabird species were reproduced with permission from Lynx Edicions and Martí Franch.

#### Comparative Genomics and Positive selection analyses

To identify genes associated with adaptation to a pelagic lifestyle in the Procellariiformes, we performed a positive selection analysis across 12 species including eight Procellariiformes species applying the HyPhy aBSREL model. We identified the hallmark of positive selection in 20 (out of the 6,172 single-copy orthologs genes), after correcting for multiple testing (Table S5). The enriched GO analysis uncovered terms related with striated muscle cell differentiation, nutrient reservoir activity, response to starvation, visual learning, positive regulation of neural retina development, olfactory receptor activity or natriuresis (Table S6). We also performed an analysis to assess the global impact of natural selection in Procellariformes (both positive and negative selection), which uncovered a total of 310 genes (Table S7). The GO terms enriched in these genes include wound healing, response to wounding, inflammatory response, sensory perception of sound, smell and chemical stimulus, neurological system process, defense response, response to stress, camera-type eye development, renal system and chloride transport among others (Table S8).

Using OrthoFinder 2.3.8, we identified 182,487 N:N orthogroups across all genes identified in the 12 analysed genomes. This data, together with the estimated ultrametric tree, was used to estimate gene gains, losses, and number of genes in the ancestral nodes using BadiRate; for the analysis we selected the Free Rates (FR) model, since it was the best fitted branch model. The analysis was conducted including all orthogroups, and the minimum number of gains and losses per branch is represented in Figure 3. Our analysis showed a tendency to gain genes in Procellariiformes (+442/-34), while the branch leading to albatrosses (Diomedeidae) showed an opposite effect, with a noticeable loss of genes (+464/-3258); the branch leading to the rest of the Procellariiformes (+379/-15) is in the line of the general behavior of the tubenoses (Table S9). Within the order, families Oceanitidae and Hydrobatidae present the same trend, with the branch leading to *H. thethys* presenting a stronger gene loss balance (+325/-966) than the branch leading to the ancestor of Oceanitidae (*O. oceanicus* and *F. grallaria* (+182/-414)).

We identified three gene families significantly expanded in the branch leading to the Procellariiformes (Table S9). These families encode zinc finger proteins (OG0000000), olfactory receptors (OG0000084) and avian histones (OG0000224).

## Discussion

### A high-quality genome assembly for the most endangered seabird in Europe

The assembly length and the GC content of the Balearic shearwater hybrid assembly presented here are similar to those reported in the seven Procellariiformes genomes released by the B10K [3]. Albeit repetitive content is remarkably higher (+33.4%) in the Balearic shearwater in comparison to the other genomes of the order, but within the range of avian genomes [26]. This difference of up to a third can be due to the fact that we included a Procellariiform (*C. borealis*) repeat library prior to running RepeatMasker, achieving a more precise library that encloses clade related repeats that are present in the genome but not found by the de novo RepeatModeler library. The genome assembly completeness (BUSCO 95.9%) is slightly higher than the obtained for other recently published bird genomes [3,29], and even higher than genome assemblies including optical mapping [30]. Despite not being a chromosome-scale assembly, contiguity is also quite high (N50 2.1 Mbp), and higher than recent avian MaSuRCA hybrid assemblies [31,32]. The retrieved proteome (21,959 protein-coding genes) is similar to previous genomes [33,34], but higher than the B10K 2020 genomes used in the comparative studies in this work (mean of 16K). This is probably due to the B10K annotation pipeline being fully based on homology, whilst we also used *de novo* prediction. The functional annotation quality in terms of genes having at least a GO term (85.9%) is comparable to recent chromosome-scale genomes [34].

The mitogenome of *P. mauretanicus* spans 19,885 bp, exhibiting the same order and the *nad6* gene duplication observed in *P. lherminieri* [27]. We did not find any *cob* duplication as it occurs in the Diomedeidae family [35]. Our result supports that *nad6* duplication could be widespread in Procellariiformes [27], and, like *cob*, could have undergone various events of deletion or addition during the diversification of the order. Nevertheless, since some of the reported duplications could be artificial [36,37], to fully identify the true number of gene duplications/deletions will require additional and specific experimental analyses.

### Heterozygosity levels and historical demography

Current level of intraspecific heterozygosity is a relevant parameter to determine the adaptive capacity of a population (or species) [38]. Since the Balearic shearwater is categorised as Critically Endangered by the IUCN, we could naively expect low heterozygosity levels in the species when compared to other Procellariiformes. However, the fossil record suggests that the Balearic shearwater had a very large population (>30.000 pairs) until the arrival of human settlers in the Balearic Islands [39], which hunted shearwaters [40] and introduced invasive mammals that also predated on them [41]. In line with Genovart, Oro, Juste, & Bertorelle, (2007) results using mtDNA markers, we observed relatively high genome-wide heterozygosity levels, suggesting that the very recent demographic decline in the species is not yet visible in its genetic diversity.

Regarding the historical demography, our PSMC’ analysis shows an increase in *N*_e_ in the Balearic shearwater from around 1 Mya to later expand to reach high population sizes, until around 150,000 ya, when it suddenly suffered a sharp decline, resulting in lower *N*_e_ values maintained until 10,000 years ago. Since current PSMC’ analysis is based on the analysis of a single genome, we could not reliably infer more recent events [28]. The Plio-Pleistocene eustatic variations resulted in a loss of neritic zones as sea level regressed [43], this may represent a loss of coastal habitat availability, which added to other oceanographic alterations (changes in ocean circulation or productivity) may have been the drivers of great population losses in marine megafauna, including seabirds. In the case of the Balearic shearwater, the PSMC’ analysis shows an abrupt decay of *N*_e_ associated with a long period of low sea level during the Penultimate Glacial Period (∼194-135 kya) which may have resulted in an important loss of neritic zones.

Here, we have observed a negative correlation between heterozygosity and body size within the Procellariiformes, where small-bodied species (*O. oceanicus, F. grallaria, H. tethys* and *P. urinatrix*) have higher heterozygosities than large-bodied species (*F. glacialis, C. borealis, P. mauretanicus* and *T. chlororhynchos*). Although controversial, contrasting heterozygosity levels between species with different body-sizes has also been reported in different species including Procellariiformes [13]. Since body-size also correlates with population size and with other life-history traits, current data does not allow to determine the biological meaning of such correlation effect [13,44].

In view of the critical population declines affecting the Balearic shearwater populations, understanding its impacts on current genetic diversity of the species and among colonies will be crucial to assess the conservation status of the Balearic shearwater. Future ongoing research, using a more powerful population genomics approach, will allow to reconstruct the most recent demographic history of the species and to test the fossil-based hypotheses of a recent loss of population due to human colonization of the island, as well as why heterozygosity values have not decayed.

### Phylogeny of Procellariiformes

The study of the evolutionary relationships within the order Procellariiformes had until recently been based mainly in the phylogenetic analyses of a single gene, the mitochondrial cytochrome b or on supertree approaches combining life history, morphological and sequence data [45–47]. However, these approaches did not show enough resolution for this group, leaving several open questions. The main points that remain contentious are: 1) which family is the sister to the rest of the Procellariiformes (Diomedeidae or Hydrobatidae), 2) which is the phylogenetic position of the diving petrels (*Pelecanoides sp*.) and whether they should be placed on their own family, 3) the monophyly of the storm petrels as well as the phylogenetic relationships among the speciose Procellariidae [48–51]. More recently, the first study to use genomic data to resolve the backbone Procellariiformes phylogeny [13] reported a well-resolved phylogeny of 51 species using 4,365 ultraconserved elements (UCEs). This phylogeny recovered the albatrosses (Diomedeidae) as the sister group to the rest of Procellariiformes, the diving petrels included within Procellariidae, and the storm petrels constituting a paraphyletic group with Oceanitidae and Hydrobatidae being two separate monophyletic groups, and Hydrobatidae as sister group of Procellariidae. Our phylogenomic results using a smaller taxon sampling but a more extensive phylogenomic dataset (of up to 6,172 genes), agrees with those of [13], supporting that these phylogenetic relationships are definitive.

### Adaptation to a pelagic lifestyle in Procellariiformes

Our selection inference uncovered 20 genes evolving under positive selection in Procellariiformes, being therefore candidates to be actively involved in the adaptation of the order to a pelagic lifestyle. Indeed, the GO’s Enrichment analysis of these genes reveals biological processes related to striated muscle cell differentiation, response to starvation, and nutrient reservoir activity, that may be related to the high energy expenditure during the vast distances they cover in the open ocean, while visual related genes could be related with underwater vision to fish and night vision [52–54]. Positive selection of genes related to natriuresis also makes sense for Procellariiformes since this biological process plays a key role to maintain the osmotic equilibrium in a sodium-rich environment like the ocean [55,56], which Procellariiformes perform thanks to the development of salt glands (modified nasal glands engaged in secretion of salts). Olfactory receptors, also found here among enriched GOs of positively selected genes, showed signature of adaptive evolution in shearwaters [57], and are crucial to Procellariiformes for navigation [58–60], partner recognition and mating [61–63], finding their own burrows [64] or foraging [65–68].

We also inferred genes with intensified natural selection in Procellariiformes, for which the GOs annotations (Table S8) are similar and coherent to those in the candidate set of genes with positive selection in all tubenoses, or, in other words, related to the adaptation of the order to a pelagic lifestyle. For example, molecular functions such as sensory perception of sound, smell and chemical stimulus, neurological system process, camera-type eye development are related with oceanic navigation. On the other hand, functions such as homeostatic process, renal system, renal response, chloride transport and regulation of ion transport point to the need of maintaining osmotic equilibrium. We also found intensified natural selection in genes participating in functions related to immune response (like inflammatory response, defense response, response to wounding, wound healing, positive regulation of phagocytosis, etc.), accompanied by a relaxation of natural selection in regulators of blood constituents, induction of bacterial agglutination, regulation of antigen processing and presentation, viral budding via host ESCRT complex or macrophage antigen processing and presentation. As Procellariiformes exposure to parasites is high [69] and their life-history traits favour parasite maintenance within populations [70], the tuning between the intensification and relaxation of natural selection in multiple biological processes and molecular functions related with immune response would have emerged following an arms race-like model. For example, as many parasites of tubenoses are blood-feeding, the intensified natural selection on the thrombin-activated receptor signaling pathway (GO:0070493), may be an evolutionary response to counter the anticoagulant activity that most blood-feeding parasites present [71].

Among the gene families expanded in the branch of Procellariiformes, the one encoding olfactory receptors is remarkable as it is coherent not only with the finding of a gene with positive selection in all Procellariiformes with the same functional annotation (g16276.t1 in the *P. mauretanicus* reference annotation, Table 4) but also with 3 olfactory receptors genes with intensification selection in the same branch (g14377.t1, g16276.t1 and g17936.t1 in *P. mauretanicus* reference annotation, Table S8). This triple evidence highlights the importance of how the adaptation to a pelagic life resulted in the enhancement of the olfactory function in Procellariiformes, as we discussed already in the paragraphs above. Moreover, similar results of positive selection in olfactory genes were obtained in *C. borealis* [57]. Physiologically, tubenoses have one of the largest olfactory bulb to brain size (OB) ratio of all birds [72]

## Conclusions

Our study highlights the utility of the hybrid assembly strategy using Illumina and ONT at recovering high quality genome assemblies, especially regarding contiguity and completeness. Comparative genomics analyses identified candidate genes under selection to have played a major role in the adaptation of the Procellariiformes to a pelagic lifestyle such as changes in sensory perception, navigation, natriuresis and physiological adaptations. Regarding the phylogeny of Procellariiformes, our results gave full support to recent genomic based hypotheses in which albatrosses (Diomedeidae) are sister to the rest of Procellariiformes, storm petrels are paraphyletic and diving petrels are included within Procellariidae. The high-quality genome presented in this work will be a great tool for future population genomic analyses, that will reveal with more precision the genetic variability of the species, its recent demographic history and the potential introgression with its sister species, the Mediterranean shearwater (*P. yelkouan*). The data obtained will be of great help in future proposals of conservation and management plans for the species.

## Methods

### Genome Assembly

We performed a *de novo* hybrid genome assembly with MaSuRCA 3.3.1 [73], using short (Illumina) and long (ONT) reads. Before the assembly step, we filtered the ONT reads with a Phred quality score (Q ≥ 5) using the NanoFilt software included in NanoPack, [74]. Paired-end Illumina reads were parsed into MaSuRCA without any preprocessing, as adapters and errors are handled by the QuORUM error corrector [75], which is part of the MaSuRCA pipeline. MaSuRCA was run applying the following parameters: fragment mean (422), fragment stdev (312) and estimated genome size (1.2 Gbp). The resulting assembly was screened for contaminants with BlobTools v1.0 [76] -x bestsumorder. Assembly completeness was assessed with BUSCO 4.0.2 [77] using the 8,338 single-copy conserved genes in aves_odb10 database [78].

### Transcriptome Assembly

We trimmed RNA-seq raw reads for adapters with BBDuk (https://sourceforge.net/projects/bbmap/) (k = 17, tpe option), and used STAR 2.7.3a [79] to map the filtered reads to the newly assembled reference genome. We obtained the transcriptome assembly with Trinity 2.8.6 [80] using the genome-guided bam mode (--genome_guided_max_intron 82945). Transcripts were clustered with CD-HIT [81,82] 4.8.1 (-c 0.98) and coding regions (CDS) were predicted with TransDecoder 5.5.0 (https://github.com/TransDecoder/).

### Mitogenome Assembly

We trimmed adapters from Illumina raw reads with BBDuk (k = 23, tpe option), before using them as input to NOVOPlasty 2.7.2 [83]. The *Puffinus lherminieri* mitogenome (MH206163.1) was used as seed using the following parameters: Genome Range (16000-24000), Insert size (422), Insert range (1.74) and Insert range strict (1.3). The annotation was performed using the MITOS WebServer [84].

### Repeat annotation

We generated a *de novo* repeat library of the genome with RepeatModeler – 1.0.11 (Smit, Hubley, & Green,) on scaffolds >100 kbp. This library was combined with all avian and ancestral consensus repeats from Dfam_Consensus-20181026 [86], RepBase-20181026 [87] and the repeat annotation of the Cory’s shearwater (*Calonectris borealis*) [3], which represents the most closely related sequenced genome. Redundancies among libraries were removed with the script ReannTE_MergeFasta.pl (https://github.com/4ureliek/ReannTE). We then ran RepeatMasker 4.0.7 [85] using the combined library as a reference, with the following parameters: -xsmall -e ncbi -s -gccalc - no_is -gff.

### Structural and Functional Annotation

We performed the structural annotation with BRAKER 2.1.2 (https://github.com/Gaius-Augustus/BRAKER) (--etpmode) using data from both the Cory’s shearwater proteome [3], and the RNA-Seq data generated in this work. Since the inclusion of RNA-Seq data appeared detrimental, we excluded this piece of information to perform the final annotation using the soft-masked genome with BRAKER 2.1.2 (--prg=gth --trainFromGth).

We made the functional annotation of the predicted genes using a similarity-based approach. We determined the protein domains with InterProScan 5.31-70.0 [88], used BLASTP [89,90] (-evalue 1e-5; -max_target_seqs 10) against the Swiss-Prot database [91] and the Cory’s shearwater and the Zebra finch reference (UP000007754) proteomes. Transcripts were annotated in the same manner. We also annotated the ncRNAs using cmscan from INFERNAL 1.1.2 [92] with the covariance models (CMs) from the Rfam 14.1 database, and tRNA genes using tRNAscan-SE 2.0.5 [93].

### Demographic History

We used MSMC2 [28] to infer the historical demography of the Balearic shearwater. MSMC2 implements a MSMC model, which allows the estimation of the effective population size (*N*_e_) over time. To generate input files for MSMC2, we mapped Illumina short reads to scaffolds larger than 1 Mbp (343 scaffolds spanning 71.8% of the assembled genome) using BWA-MEM 0.7.17 [94], as recommended in [95]. First, we called the SNPs using samtools mpileup (Samtools mpileup 1.9 -q 20 -Q 20 -C 50) and then bcftools 1.9 -c -V indels. The input files were then generated by converting the SNPs obtained to MSMC input format using the bamCaller.py script accounting for the mean coverage of each scaffold. Multiple sequentially Markovian coalescent (MSMC) for two haplotypes, known as PSMC’, was run with MSMC2 with time patterning specified as -p 1*4+30*2+1*4+1*6+1*10.

We ran 100 bootstraps of 29 pseudo-chromosomes [96] sampling 20 chunks of 1.508.752 bp with replacement using multihetsep_bootstrap.py. We scaled time and population size using a generation time for the Balearic shearwater of 12.8 years [15] and the Northern fulmar (*Fulmarus glacialis*) mutation rate (2.89×10^−9^ substitutions per nucleotide per generation, [2].

### Genome-wide heterozygosity

We estimated genome-wide heterozygosities using information of a single individual from all eight Procellariiformes species studied. We applied the [97] method, with minor modifications to take genome fragmentation into consideration, since we included genome assemblies with varying amounts of contiguity. The DNA sequence data (genome assemblies and whole-genome sequencing data) were downloaded from NCBI (PRJNA261828, PRJNA545868; [1,3]. For each species, adapter-trimmed reads were aligned to its genome assembly using bwa mem [98], bam files were merged using Picard-Tools (http://broadinstitute.github.io/picard/) and variants were called using the GATK 4.1.9 HaplotypeCaller and GenotypeGVCFs [99]. Sites with a coverage <1/3X or >2X of the average coverage depth (of the particular genome) were filtered out using VCFtools 0.1.15 [100]. We computed per-site heterozygosity as the proportion of heterozygous sites per total number of called genotypes within a single individual in nonoverlapping 25Kb windows across each scaffold. Windows with less than 50% of net sites (those excluding missing or filtered sites), were excluded from the analysis.

### Orthology inference

We performed the phylogenomic and comparative genomics analyses including information from 12 species with an available genome assembly: eight Procellariiformes (*P. mauretanicus, Thalassarche chlororhynchos, Hydrobates tethys, Oceanites oceanicus, Fregetta grallaria, Pelecanoides urinatrix, Fulmarus glacialis* and *Calonectris borealis*), and four outgroups, *Aptenodytes forsteri, Pygoscelis adeliae* (Sphenisciformes); *Egretta garzetta, Phalacrocorax carbo* (Pelecaniformes). We inferred orthologous genes across the proteomes of these 12 species using OrthoFinder 2.3.8 [101] with default parameters.

### Phylogenetic relationships

We built a multiple sequence alignment (MSA) for each 1:1 orthologs with PRANK v.100802 [102], using both coding sequences (CDS) (-codon -noxml -notree -F) and amino acid sequences (-noxml -notree -F). Individual alignments were concatenated with catfasta2phyml v1.1.0 (https://github.com/nylander/catfasta2phyml) to create a CDS supermatrix and an amino acid supermatrix. Only locus with data for all the twelve species have been considered. Fourfold degenerate sites (4D) for the CDS supermatrix were extracted with MEGA X [103]. We performed unpartitioned maximum likelihood (ML) phylogenetic analyses using IQ-TREE 1.6.12 [104] (-bb 1000) both for 4D and amino acid supermatrices. Optimal models of sequence evolution were obtained with ModelFinder [105] according to Bayesian information criterion (BIC), and the resulting best-fit models were GTR+F+R2 for 4D and HIVb+F+R3 for amino acid supermatrix. Node support was obtained with Ultrafast Bootstrap [106].

To explicitly account for incomplete lineage sorting (ILS) under the Multispecies Coalescent Model (MSC), we inferred the species tree using the summary coalescent approach, as implemented in ASTRAL-III 5.6.3 [107]. We first obtained all gene trees (for each 1:1 orthologous genes) using IQ-TREE 1.6.12, and inferred the species tree and its normalized score (from both CDS and amino acid gene trees) using ASTRAL-III. We generated an ultrametric tree with r8s v.1.81 [108] using the 4D supermatrix ML tree. We used four calibration points (in myr): root (max_age=84 min_age=73, [109]), most recent common ancestor (MRCA) of Spheniscidae (min_age=12.6, [110]), MRCA Procellariiformes (min_age=49, [111]), MRCA Procellariidae (min_age=14, [112]), retrieved from TimeTree [113]. We used the penalized likelihood (PL) method and the Truncated Newton (TN) algorithm, smoothing parameter was set to 100.

### Positive Selection Analysis

We evaluated the selective constraints of genes that could be associated with pelagic lifestyle. For this purpose, we performed the analysis with HyPhy 2.5 [114], using Procellariiformes data (1:1 MSAs). Prior to the analysis, non-reliable positions across all 1:1 orthologs MSAs were filtered with ZORRO [115] (default options; MSA with average quality < 5 were filtered). We used the aBSREL method [116] to test for positive selection, and the RELAX method [117] to test for relaxed/intensified selection. We also performed a Gene ontology (GO) enrichment analysis of the candidate genes using the GOstats [118] R package against the background GOs of 1:1 orthologs.

### Gene family evolution

We estimated gene turnover rates, the number of gene gains and losses across the phylogeny lineages, and inferred gene family contractions and expansions using BadiRate 1.7 [119]. For the analysis we first inferred the orthogroups with OrthoFinder 2.3.8, and we used the calibrated ultrametric tree estimated with r8s. We tested, under the Birth-Death-Innovation (BDI) model for turnover rates, several biological relevant hypotheses with three different branch models: Free Rates (FR), Global Rates (GR) and Branch-specific Rates (BR), and chose the best model based on the lowest AIC value. To ensure an appropriate convergence we ran multiple times each model.

## Acknowledgements

We are grateful to David García and Maite Louzao for kindly providing samples and the Govern Illes Balears for research permits (CEP19/2019). This research was supported by Fundación Banco Bilbao Vizcaya (Spain), Project 062-17; by the Ministerio de Economía y Competitividad of Spain, projects CGL2016-78530-R, PGC2018-093924-B-100 and PID2019-103947GB.

## Author contributions

JFO, JGS, MR and JR, conceived the study. CCC, JFO, MG and JGS, performed samplings. CCC, JFO performed wet lab work. CCC, JFO and JV performed the bioinformatic analyses. CCC, JFO, MR and JR, interpreted the genomic data. MG, JGS discussed on biological data. CCC, JFO, MR and JR drafted the first version of the manuscript. All authors revised and approved the final version of the manuscript.

## Data Availability

The whole-genome shotgun project has been deposited at DDBJ/ENA/GenBank under the Bioproject PRJNA780920, and BioSamples SAMN24039388, SAMN23492024 and SAMN23212142. The raw reads are also available in the Sequence Read Archive (SRA) under the Bioproject accession. Other relevant datasets, such as those including the structural and functional annotations, are available in https://github.com/molevol-ub/Puffinus_mauretanicus_genome, and in the Supplementary Information online.

