## Supplementary Figures for "The genome of the Balearic shearwater (*Puffinus mauretanicus*), a Critically Endangered seabird: a valuable resource for evolutionary and conservation genomics"

### Supplementary Tables

Table S1. Summary of the RepeatMasker analysis results.

Table S2. Annotated predicted protein-coding genes, their lengths and their functional annotation based on matches against the curated databases.

Table S3. Annotated predicted transcripts, their lengths and their functional annotation based on matches against the curated databases.

Table S4. Mean coverage and standardized coverage (gene coverage/whole mitogenome coverage) for each annotated mitochondrial gene.

Table S5. Genes with evidence of positive selection, and their functional annotation, in the Procellariiformes included in the comparative genomics study.

Table S6. GOs terms enriched in Procellariiformes dataset (positive selection) with its description and *p*-values. Dispensable GOs are collapsed.

Table S7. RELAX results: list of genes under selection (both relaxed or intensified) in Procellariiformes.

Table S8. GO terms enriched in the Procellariiformes (from RELAX dataset; Table S7). Dispensable GOs are collapsed.

Table S9. Outlier Families in the Procellariiformes branch (BadiRate analysis), with its *p*-value, FDR_P-value, and its functional annotation.

### Supplementary Figures

Figure S1. Annotated mitogenome of *P. mauretanicus*.

Figure S2. Species tree made with CDS supermatrix 4D positions with IQ-TREE.

Figure S3. Species tree made with amino acid supermatrix with IQ-TREE.

Figure S4. Species tree made with ASTRAL using CDS gene tree quartets.

Figure S5. Species tree made with ASTRAL using amino acid gene tree quartets.

### Other Supplementary Material

Other relevant datasets, such as those including the structural and functional annotations, are available in:

https://github.com/molevol-ub/Puffinus_mauretanicus_genome

Figure S1. Annotated mitogenome of *P. mauretanicus*.

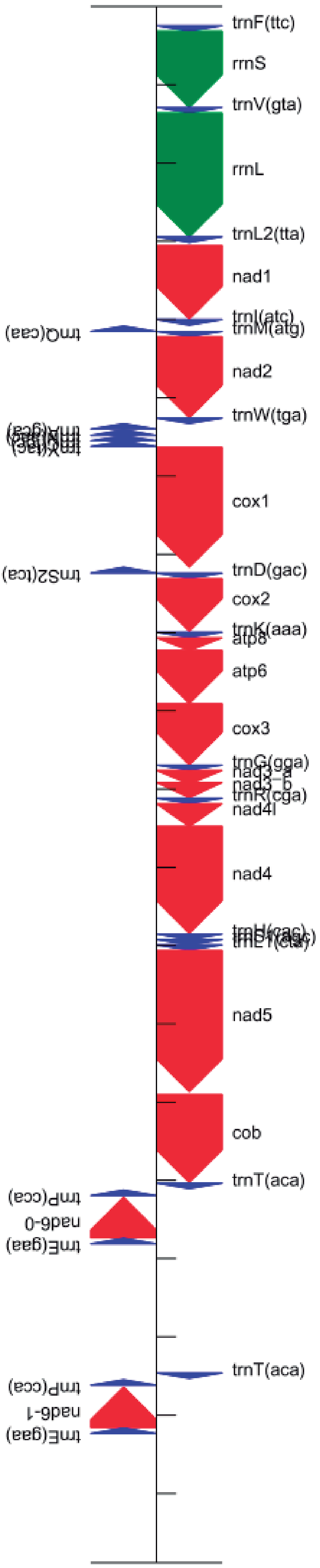

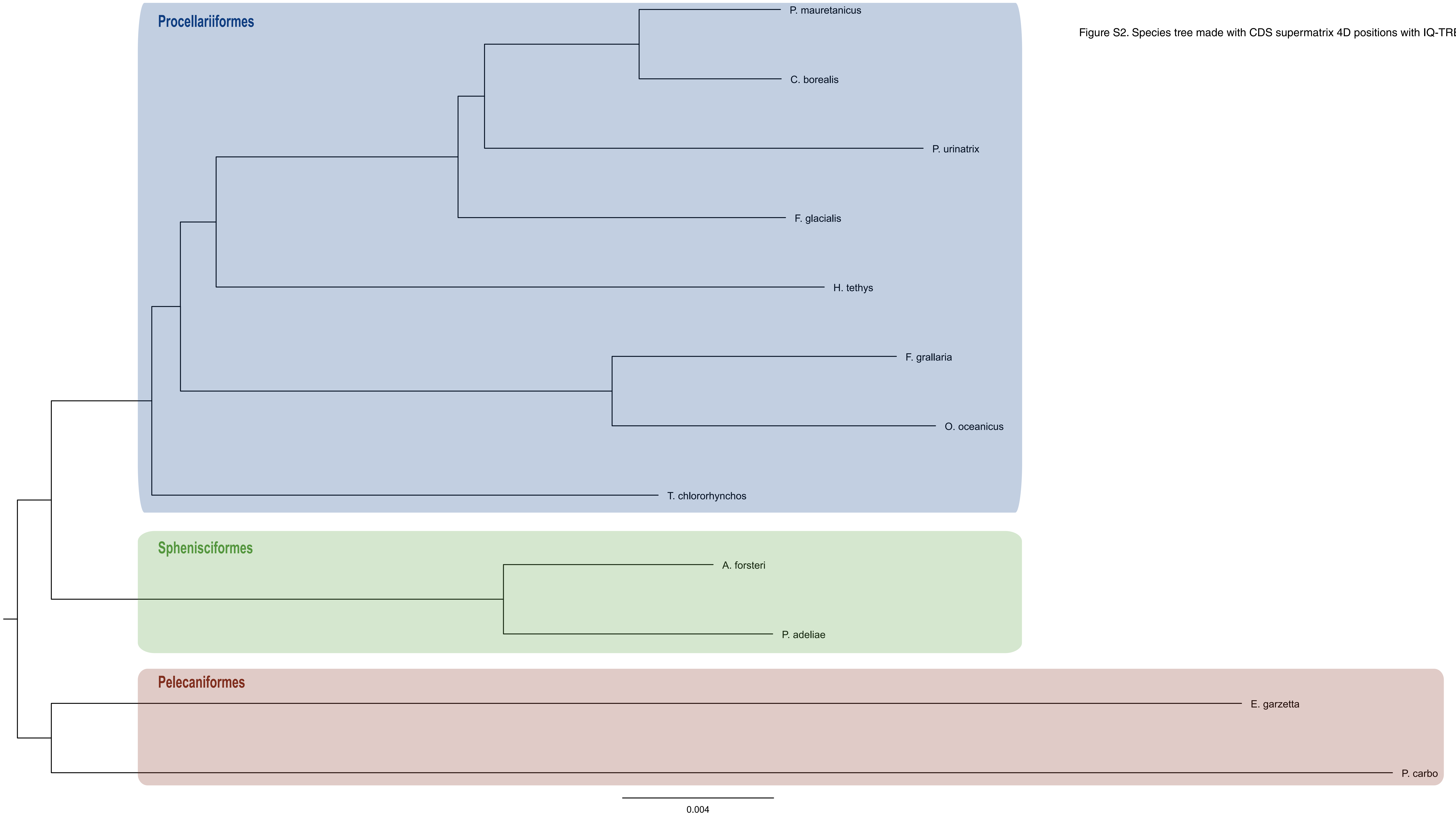


Figure S2. Species tree made with CDS supermatrix 4D positions with IQ-TREE.

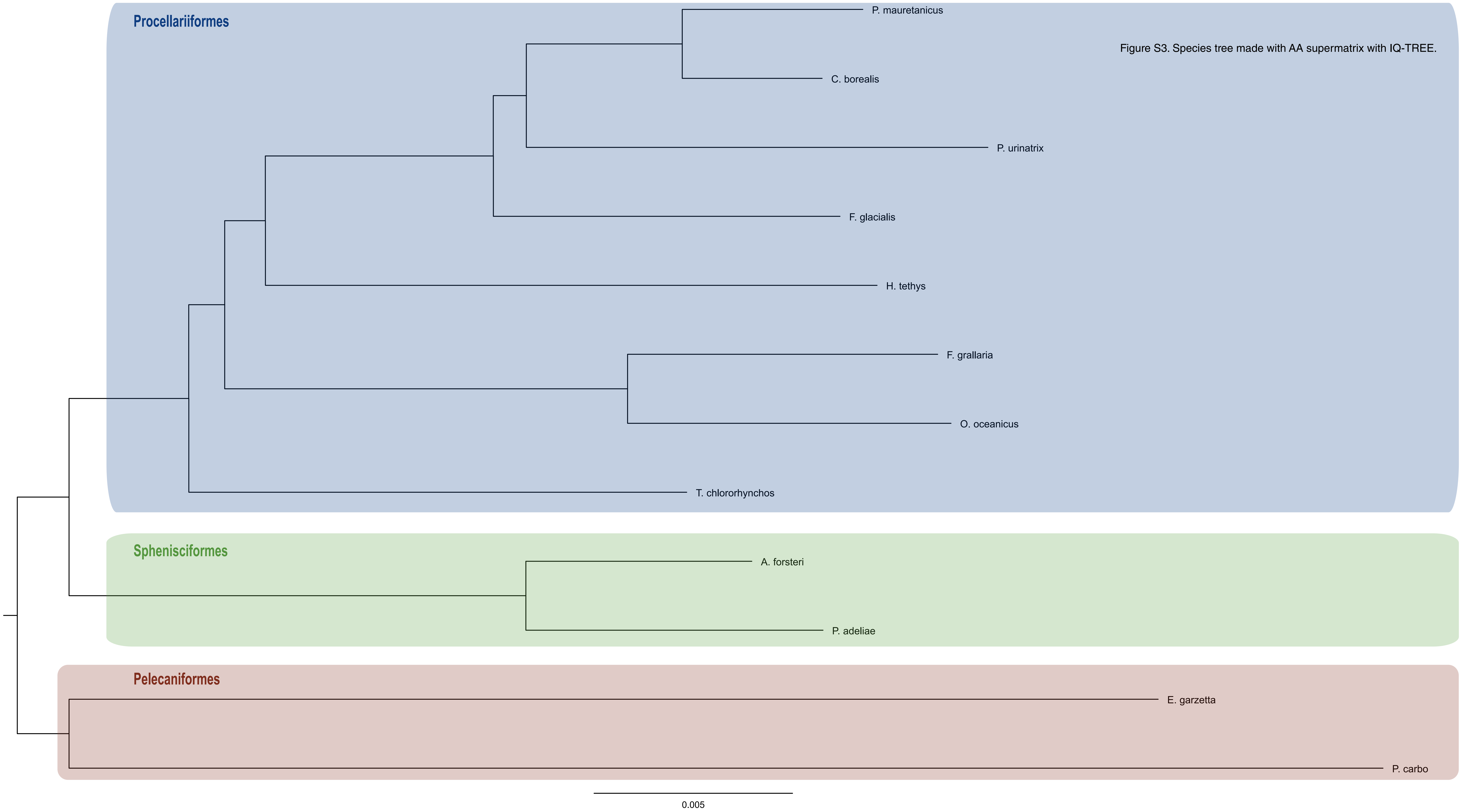


Figure S3. Species tree made with AA supermatrix with IQ-TREE.

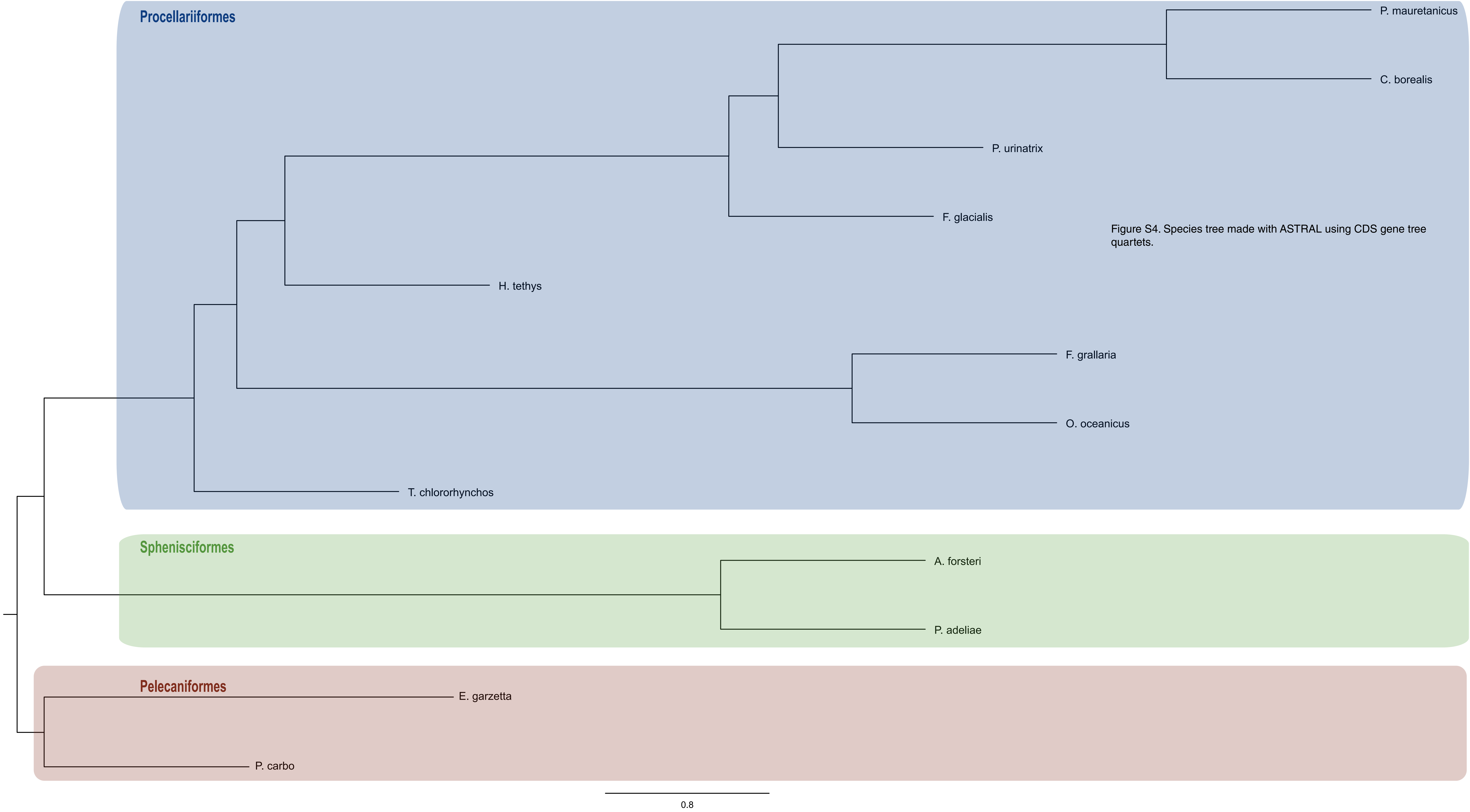


Figure S4. Species tree made with ASTRAL using CDS gene tree quartets.

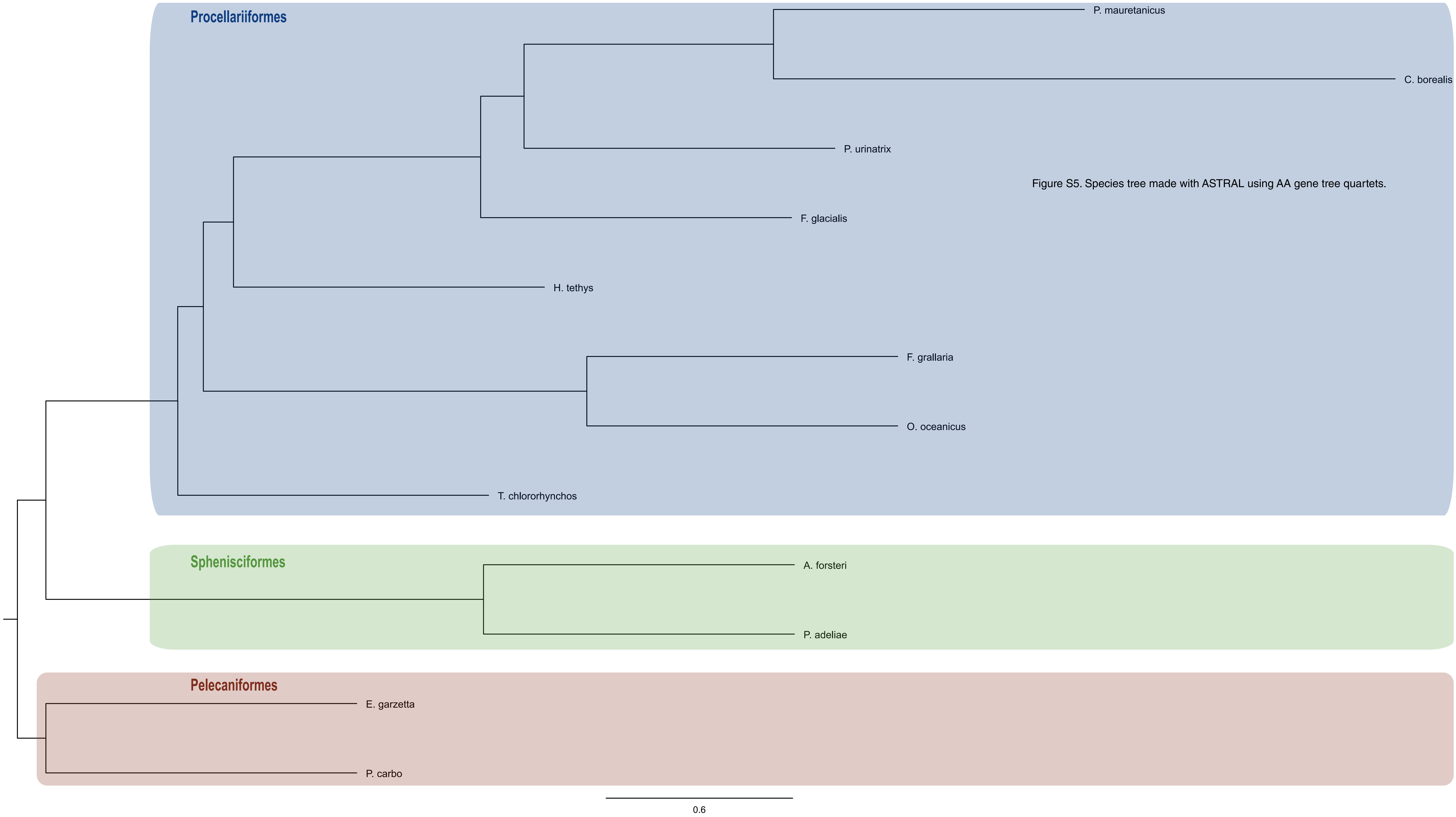


Figure S5. Species tree made with ASTRAL using AA gene tree quartets.
